# Speedy-PASEF: Analytical flow rate chromatography and trapped ion mobility for deep high-throughput proteomics

**DOI:** 10.1101/2023.02.17.528968

**Authors:** Lukasz Szyrwiel, Christoph Gille, Michael Mülleder, Vadim Demichev, Markus Ralser

## Abstract

Increased throughput in proteomic experiments can improve accessibility of proteomic platforms, reduce costs and facilitate new approaches in systems biology and biomedical research. Here we propose Speedy-PASEF, a combination of analytical flow rate chromatography with ion mobility separation of peptide ions, data-independent acquisition and data analysis with the DIA-NN software suite, for conducting fast, high-quality proteomic experiments that require only moderate sample amounts. For instance, using a 500-μl/min flow rate and a 3-minute chromatographic gradient, Speedy-PASEF quantified 5,211 proteins from 2 μg of a mammalian cell-line standard at high quantitative accuracy and precision. We further used Speedy-PASEF to analyze blood plasma samples from a cohort of COVID-19 inpatients, using a 3-minute chromatographic gradient and alternating column regeneration on a dual pump system, for processing 398 samples per day. Speedy-PASEF delivered a comprehensive view of the COVID-19 plasma proteome, allowing classification of the patients according to disease severity and revealing plasma biomarker candidates. Speedy-PASEF thus facilitates acquisition of high-quality proteomes in large numbers.

## Introduction

Recently, significant efforts have been directed towards making proteomic experiments faster and more accessible. Fast proteomics facilitates new applications, such as the analysis of large human patient cohorts for biomarker discovery as well as for precision medicine, time-resolved profiling of multiple conditions in systems biology or discovery approaches such as the screening of drug panels and genetic perturbations^1–4^. In the past few years, this increasing demand for fast proteomic workflows has led to the establishment of automated sample-preparation methods, LC-MS approaches with fast chromatographic gradients and higher chromatographic flow rates, faster mass spectrometers, as well as new data-independent acquisition (DIA) and data processing methods, allowing proteomic profiling of hundreds of samples per day on a single mass spectrometer.

Notably, some of the technological developments that led to higher throughput have also improved the consistency of protein identification, protein quantification and reduced batch effects, rendering proteomics increasingly routine-compatible^5–17^. Further, as both instrument time and hands-on time are key cost factors in mass spectrometry based proteomics, the fast sample throughput has rendered proteomic experiments more economical.

Recently, notable progress in accelerating throughput has been achieved with the introduction of analytical flow rate chromatography, with flow rates of several hundred microlitres per minute, in combination with DIA proteomic experiments. Analytical flow rate chromatography reduces sample carry-over, accelerates column equilibration, and allows fast chromatographic gradients. Moreover, it is robust when applied to large sample series^5,6,9^. It results in sharper peptide elution and hence higher peak capacity, leading to cleaner spectra, this being particularly beneficial for fast chromatographic gradients^5^. However, analytical flow rate proteomics faces the challenge of a high degree of sample dilution. Therefore, it requires high sample amounts, and often reduces proteomic depth – important limitations for a range of applications.

Trapped ion mobility separation (TIMS) implemented in timsTOF instruments (Bruker) with the dia-PASEF technology has been shown to improve proteomic analyses, including in combination with fast gradients, by both increasing the sensitivity and adding another dimension of peptide separation^7,8,18^. Here we introduce a proteomic platform, termed Speedy-PASEF, that involves a combination of analytical flow rate chromatography, dia-PASEF, and analysis with the neural-network-enabled DIA-NN software^15^, to create a robust and sensitive workflow for high-throughput proteomics. Speedy-PASEF reaches considerable proteomic depth despite moderate sample requirements. For instance, it quantified 5.8 thousand proteins in single 2-μg injections of a K562 tryptic cell-line standard using a 5-minute active gradient and 500-μL/min chromatography at high quantitative accuracy and precision. Further, drawing upon the high sensitivity of the DIA-NN data processing software, as well as the sensitivity increase driven by TIMS, our workflow quantified 2,038 proteins from merely 25 ng of a mammalian cell line standard, while maintaining the robustness of analytical flow rate separation. Applied to measure the plasma proteomes from a cohort of COVID-19 patients, we demonstrate the utility of our workflow for high-throughput clinical proteomic applications in conjunction with a dual-pump system for alternative column regeneration, facilitating the analysis of hundreds of samples per day.

## Results

### Instrument setup and settings

The high-throughput proteomic workflow was established using a timsTOF Pro mass spectrometer (Bruker) coupled to an analytical flow liquid chromatography (LC) system (Infinity 1290 II; Agilent), equipped with either one or two columns for chromatographic separation. The one-column system is, at least in theory, more robust, and more broadly accessible. Instead, the two-column setup that uses an LC system with two binary pumps allows for concomitant peptide separation on one column during the equilibration of the other column, thus reducing the total run time, as one column equilibrates whilst the other column separates.

Our liquid chromatography methodology was derived from a method reported previously that used an 800-μl/min gradient and a 5-minute water-to-acetonitrile gradient on a C18 ZORBAX Rapid Resolution High Definition (RRHD) 50 × 2.1 mm, 1.8 μm column operated at 30°C^5^. Here we used a shorter column (30 × 2.1 mm Luna OMEGA 1.6 μm C18 100 Å; Phenomenex) and also increased the column temperature to 60°C, which led to better separation of peptides and resulted in about 10% increase in peptide identifications. The one-column setup was further optimized for sensitivity by reducing the flow rate from 800 to 500 μl/min. The dual-column setup was optimized for separation efficiency at the expense of sensitivity, using a 850-μl/min flow rate, to target applications in which the sample is available in virtually unlimited amounts – such as most plasma and serum proteomic workflows. The resulting throughput of the system was 8.6 minutes per injection (corresponding to 168 samples/day) for a 5-minute active gradient on a one-column setup, 6.1 minutes per injection (236 samples/day) for a 3-minute active gradient on a one-column setup, and 3.7 minutes per injection (398 samples/day) for a 3-minute gradient on a two-column setup.

We tested two ion-source options, one known as the ESI source (Bruker) and a vacuum-insulated probe-heated ESI (VIP-HESI; Bruker). Application of heated electrospray over the conventional ESI source supports vaporization at the maintained temperature to increase desolvation of the analyte ions^19^. We achieved a better proteomic depth with the VIP-HESI source. We further investigated changes in transfer time in the range of 55–65 μs and in the collision energy maximum value (up to 80 eV), but did not observe a significant effect on the number of peptide identifications and thus used the default values of 60 μs and 59 eV.

The dia-PASEF^8^ acquisition scheme was optimized for a cycle time estimate of 0.7s, maximizing the sensitivity of the TOF acquisition while maintaining the required number of data points per peak at a level of ∼2.5 – 3 data points at full width half maximum on average, similar to our previous work^5^. The dia-PASEF MS/MS isolation windows scheme (Supplementary Figure S1) was selected in a fashion similar to previous works^8,18^ to cover most of the charge 2 precursor ions in the range m/z 401–1226 and 1/K_0_ 0.72–1.29, using 33 × 25 Th windows, with accumulation and ramp times of 72 ms. The mass spectrometer was operated in the ‘high sensitivity detection’ (‘low sample amount’) mode.

### Identification performance

To establish the protein identification performance and quantitative reproducibility of our method, we acquired a dilution series of a commercial tryptic digest standard of a human cell line (K562, Promega; Methods), injecting amounts ranging from 10 ng to 3,000 ng in triplicates, using a 5-minute gradient at a 500-μl/min flow rate. The data were processed using DIA-NN 1.8.1^15^, which contains an ion mobility module optimized for dia-PASEF^7^. For this, we created a spectral library consisting of 104,026 peptide precursors (i.e. peptides with a specific charge) by using spin-column-based high-pH separation of the sample in 10 fractions, followed by 20-minute gradient analysis of each fraction in dia-PASEF mode, using the same LC-MS setup and data processing using DIA-NN (Methods). The data were filtered at 1% run-specific precursor q-values and, for protein-level analysis, 1% run-specific protein q-values, to ensure strict false discovery rate (FDR) control for individual acquisitions. We note that the use of global protein FDR control instead would have resulted in higher data completeness.

The identification numbers ranged from 1,253 proteins and 6,740 peptide precursors for 10 ng to 5,844 proteins and 67,098 precursors for 3,000 ng (Figure 1a). High quantitative precision was maintained, with median protein coefficients of variation ranging from 17% (10 ng) to 5% (3,000 ng).

**Figure 1.**
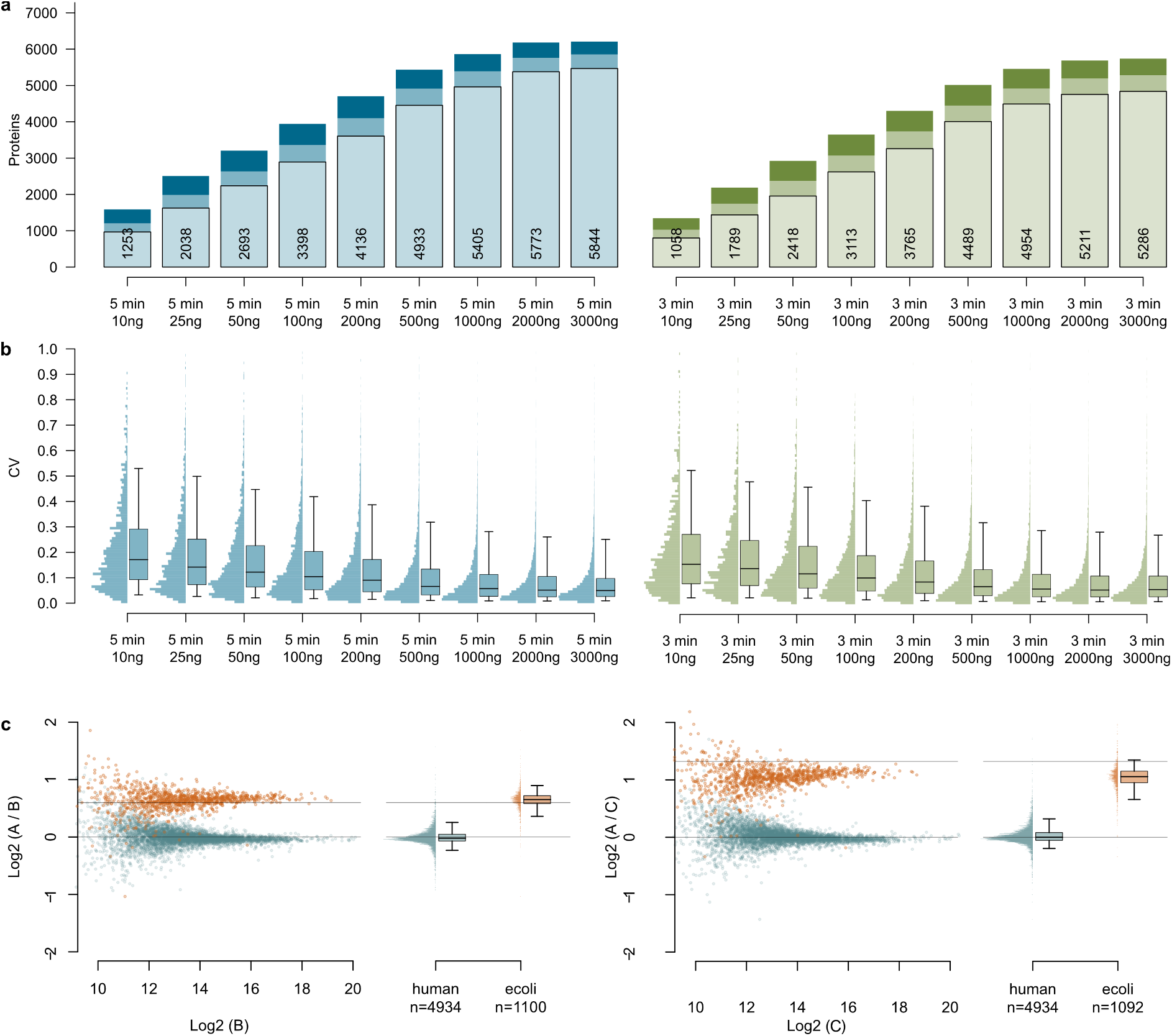
Performance characteristics of Speedy-PASEF using analytical flow rate chromatography. **a** Numbers of quantified proteins for different injection amounts of a K562 tryptic digest analyzed with a 5-minute (left) or 3-minute (right) analytical flow gradient (500 μl/min). Proteins detected in 1, 2, or all 3 injection replicates for each amount are shown with different color shades; average numbers are indicated. **b** The respective coefficient of variation (CV) distributions. The boxes correspond to the interquartile range, with the median indicated, and the whiskers extend to the 5–95% percentiles. **c** Protein ratios between different mixtures of human (K562) and *E. coli* tryptic digests, measured using a 5-minute analytical flow gradient (500 μl/min). Left: mixtures A and B (human A:B = 1:1, *E. coli* A:B = 50:33). Right: mixtures A and C (human A:C = 1:1, *E. coli* A:C = 50:20).

We further investigated the performance of Speedy-PASEF when operated with a 3-minute active gradient (Figure 1ab, right panels). Here, the identification numbers ranged from 1,058 proteins and 5,591 peptide precursors for 10 ng to 5,286 proteins, and 53,207 precursors for 3,000 ng injected standard. High quantitative precision was maintained, with median protein coefficients of variation ranging from 16% (10 ng) to 6% (3,000 ng). Hence, the faster method demonstrated a proteomic depth close to that of the longer method, combined with comparable quantitative precision, making it an attractive choice for high-throughput experiments.

### Speedy-PASEF maintains quantitative accuracy

While analytical flow rate chromatography has beneficial effects on peak capacity and method robustness^5^, fast gradients in general limit peptide separation in comparison to long-gradient chromatography, resulting in more complex convoluted spectra and thus potentially affecting quantitative accuracy. To validate the quantitative performance of Speedy-PASEF, we followed the LFQBench-type approach as established by Navarro and colleagues^20^. LFQbench evaluates the accuracy of proteomic experiments on the basis of their ability to correctly recover the known protein ratios between mixtures of peptide preparations from different organisms. Here we spiked in an *E. coli* tryptic digest standard (Waters, 500, 333, or 200 ng per injection) in the K562 standard (2000 ng per injection) and analyzed the samples in triplicates using 5-minute-gradient Speedy-PASEF and the library-free processing mode in DIA-NN (Figure 1c). We obtained 4,934 human protein ratios and 1,100 (between 500 ng and 333 ng) or 1,092 (between 500 ng and 200 ng) *E. coli* protein ratios between the mixtures. To put these numbers into context, the 120-minute gradient data acquired as part of the original LFQbench study using nano-flow rate chromatography and a mass spectrometer without a TIMS device^20^ yielded 4,636 proteins cumulatively identified across three species (human, yeast, *E. coli)* when processed using the OpenSWATH pipeline^21^. The analysis of the Speedy-PASEF data further revealed comprehensive precision and accuracy of quantification, with the distributions of human and *E. coli* protein ratios well separated and clustered close to the expected values. For both mixture comparisons (*E. coli* 500 ng : 333 ng, 500 ng : 200 ng) the median coefficients of variation for human proteins were 7%. The standard deviations of log _2_ ratios between mixtures were 0.16 (human), 0.19 (*E. coli*) for the *E. coli* 500 ng : 333 ng comparison and 0.18 (human), 0.23 (*E. coli*) for the *E. coli* 500 ng : 200 ng comparison.

### Comprehensive profiling of neat human plasma of a COVID-19 inpatient cohort

One key application of fast proteomic experiments is plasma or serum proteomics. Plasma proteomics drives translational research, as it provides information about disease phenotypes, identifying novel biomarkers and predicting disease progression, complementary to established diagnostic tests^22,23^. In plasma proteomics, throughput and robustness of the workflow are of particular relevance, allowing to scale up the studies, leading to statistical confidence and better performance of machine learning models^1,5,6,23–28^. Moreover, human blood plasma is a matrix that differs quite substantially from the one of a cellular protein extract. Most importantly, some ∼300 proteins constitute > 99,99% of the protein mass in plasma, but these proteins are present in many iso- and proteoforms ^29^. In order to fully exploit the throughput of Speedy-PASEF, we used a dual pump and dual column configuration of the LC system. This configuration accelerates sample throughput to about 400 samples per day, as one column can equilibrate while the other column separates. Moreover, this setup allows for long column equilibration and extensive column washing despite the fast throughput, which is beneficial for chromatographic stability, decreases the degree of carryover, and prolongs the column lifetime.

To evaluate the performance of this Speedy-PASEF setup for undepleted plasma proteomics (in this case citrate plasma), we studied a cohort consisting of samples derived from 30 COVID-19 inpatients with different disease severity grades, as well as from 15 healthy donors, described in our previous work^5^. Including the control samples, the plasma cohort (73 injections) was processed in less than 4.5 hours of instrument time (Figure 2a).

**Figure 2.**
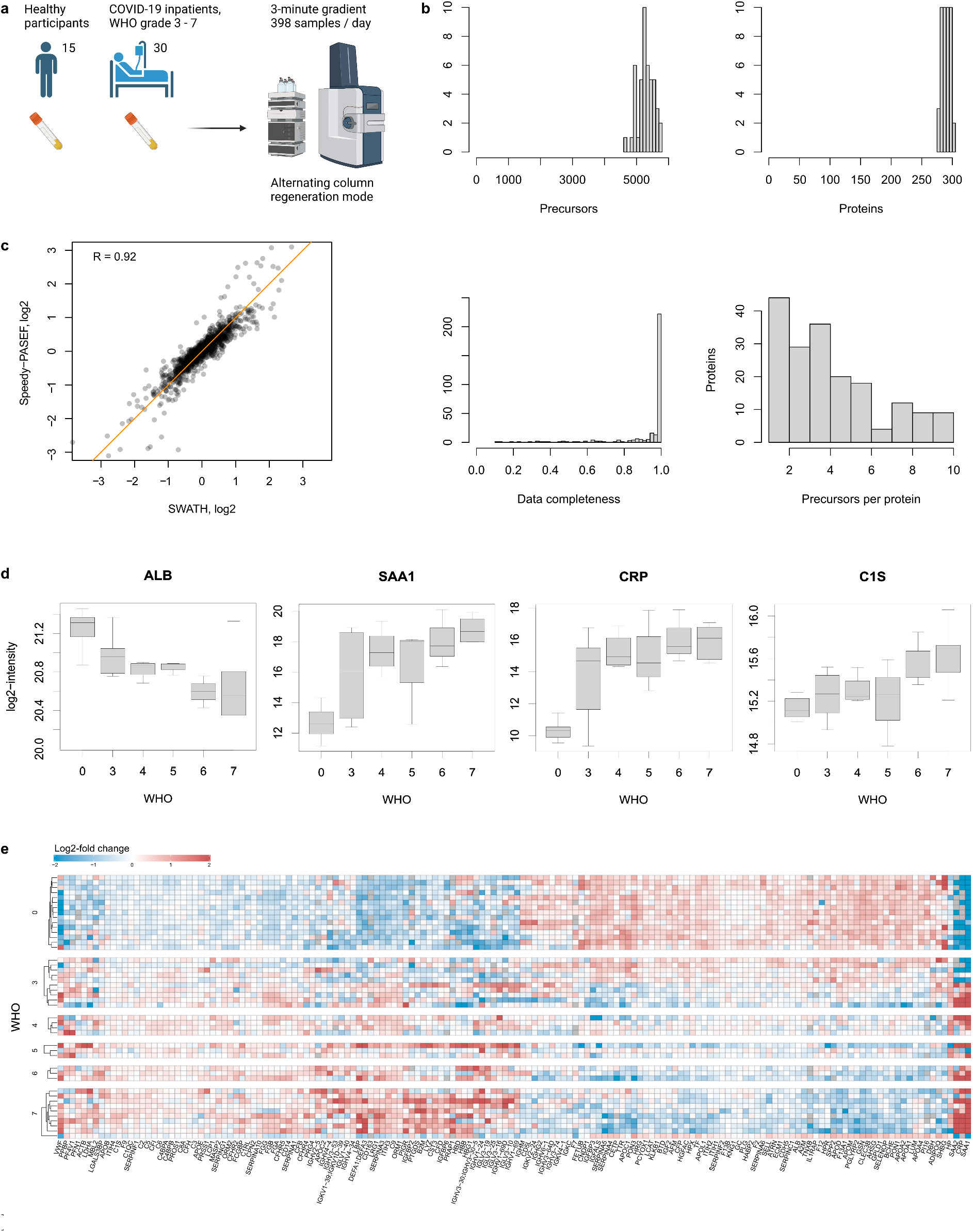
Plasma proteomics of a COVID-19 inpatient cohort using Speedy-PASEF. **a** Plasma samples from 30 COVID-19 inpatients with disease severity grades ranging from mild (WHO ordinal scale for clinical improvement (‘WHO grade’) 3 to critical (WHO grade 7), as well as from 15 healthy donors (WHO grade 0), were analyzed using 3-minute-gradient Speedy-PASEF. Data acquisition was completed in less than 4.5 hours of instrument time. **b** Numbers of precursors (top left) and proteins (top right) as well protein data completeness (bottom left) and numbers of precursors per protein (bottom right) given as histograms. **c** Comparison of the centered log_2_ levels of 25 proteins (A1BG, ALB, APOA1, APOC1, C1R, C1S, C2, C8A, CD14, CFB, CFH, CFI, FGA, FGB, FGG, GSN, HP, ITH3, ITIH4, LBP, LGALS3BP, LRG1, SAA1, SERPINA10, TF) which we previously found with high confidence to be differentially abundant depending on COVID-19 disease severity based on SWATH proteomic measurements^5^, between SWATH (5-minute gradient, triplicate injections) and Speedy-PASEF (3-minute gradient, single injections). **d** Variation of selected gene products depending on disease status and severity; boxes correspond to the interquartile range, medians are indicated, whiskers extend by up to 1.5-times the interquartile range from the box. **e** The relative log_2_ intensities of 156 proteins that are differentially abundant depending on disease status or severity grade (FDR <= 0.05), grouped by the severity grade. Each row corresponds to a single sample.

The data were analyzed with the same public spectral library derived from the DioGenes study ^24^ that we used previously in the analysis of the COVID-19 cohort proteomes^5^. Speedy-PASEF demonstrated highly robust performance, identifying and quantifying on average 5,286 peptide precursors and 290 proteins per sample, with 94% data completeness at the protein level and a median of 4 precursors identified per protein (Figure 2b). A non-parametric differential abundance analysis (Methods) revealed the association of 156 proteins (5% FDR) with disease status or severity (Figure 2de). Consistent with previous studies, proteins identified to be differentially abundant depending on COVID-19 severity reflect known biological processes activated by the disease, including components of the innate immune system (e.g. complement system), coagulation factors, and apolipoproteins^1,25,30^. For example, we observed a consistent signature of the acute phase response induced by COVID-19, including downregulation of plasma albumin (ALB), as well as upregulation of C-reactive protein (CRP) and serum amyloid A 1 and 2 (SAA1 and SAA2). The upregulation of multiple complement proteins was also observed, as well as modulation of several apolipoproteins, including upregulation of APOB and downregulation of APOA1, APOA2, and APOC1 (Figure 2de).

We further compared these results recorded with Speedy-PASEF to those we previously obtained by measuring the same cohort with SWATH^31^ using a 5-minute 800-μl/minute analytical flow active gradient, on an previous generation of a fast qTOF mass spectrometer (Triple TOF6600, Sciex) ^5^. In total, 256 proteins were detected by both technologies. However, Speedy-PASEF achieved a 96% data completeness for these proteins, as opposed to 89% data completeness by SWATH on a TripleTOF6600 instrument, despite the previous experiment featuring a longer gradient and triplicate injections per sample. Comparing the centered log_2_ intensities of the 25 proteins that we previously found with high confidence to be indicative of disease status^5^ between the technologies, we observed high correlation (R = 0.92) as well as similar dynamic range (Figure 2c). Thus, on the plasma samples, we obtain highly congruent quantitative results with Speedy-PASEF in comparison to our previous study that used a different high-throughput proteomic strategy.

## Discussion

We demonstrate that the combination of analytical flow rate chromatography and dia-PASEF generates a quantitatively reliable proteomic platform for high-throughput studies while attaining a comprehensive proteomic depth. Speedy-PASEF draws upon the high chromatographic peak capacity of analytical flow rate chromatography^5^ as well as the precursor ion separation in the TIMS dimension^32^ to achieve low levels of interferences in the resulting MS/MS spectra and thus promote high identification rates and reliable quantification using DIA acquisition^7,8^. For instance, we report the identification of 5,844 proteins in single injections of 3,000 ng K562 tryptic digest analyzed with a 5-minute gradient, which is, to our knowledge, the highest proteomic depth reported for methods of similar throughput. The workflow also shows high reproducibility, with median protein CV values of 7% for microgram-range injection amounts, and maintains quantitative accuracy. Speedy-PASEF also yielded 5,286 proteins identified with a 3-minute gradient, comparable to previous reports for a 5.6-minute gradient on a microflow setup^7^.

The TIMS separation in dia-PASEF has a further benefit of boosting the duty cycle of MS/MS acquisition, that is the proportion of ionized peptides that are being fragmented, and thus promoting the sensitivity of peptide detection^8^. Indeed, we could identify and reproducibly quantify 3,113 (3 minutes) or 3,398 (5 minutes) proteins from 100 ng injections. To put these results in perspective, the proteomic depth reported here for 100 ng injections acquired with a 3-minute gradient exceeds the numbers recently obtained on timsTOF Pro with a 5.6-minute microflow method from 200 ng, using the data processing software that was state of the art at the time^8^.

When applied to plasma proteomics, using the two pump system and alternating column regeneration mode, Speedy-PASEF offers high throughput while maintaining comprehensive proteomic depth and achieving high data completeness, making it a promising technology for large-scale profiling of proteomic samples, for example, to conduct plasma or serum cohort studies. Within a short measurement time, Speedy-PASEF identified proteomic changes associated with COVID-19 and enabled patient classification. For this application, the quantitative results we obtained with Speedy-PASEF were in high agreement with our previous study on the same cohort acquired with SWATH-MS^5^. Speedy-PASEF hence expands the compendium of fast proteomic methods such as Scanning SWATH or Zeno-SWATH^6,9^, a compendium that provides high-quality proteomes in combination with analytical flow rate chromatography and that has its strengths in large-scale and routine applications due to its fast throughput, high reproducibility, and robustness.

In summary, we propose Speedy-PASEF, a workflow that facilitates high-performance, high-throughput measurement of samples that are available in moderate-to-high amounts, by combining analytical flow rate chromatography^5^, dia-PASEF^8^, and DIA-NN analysis^7^. The inherent limitation of analytical flow – the requirement of high injection amounts – is alleviated by the sensitivity increase afforded by the TIMS device, extending the applicability of Speedy-PASEF to sub-100 ng sample amounts, while maintaining the robustness and peak capacity of the analytical flow rate LC platform.

## Methods

### LC-MS

We used an 1290 Infinity II chromatographic system (Agilent) coupled to a timsTOF Pro (Bruker) mass spectrometer equipped with a VIP-HESI source (3,000 V of capillary voltage, 10.0 l/min of dry gas at 280°C, probe gas flow 4.8 l/min at 450°C). The peptide separation was performed on a 30 × 2.1 mm Luna OMEGA 1.6 μm C18 100 Å (Phenomenex) column at 60°C. The 5-minute one-column active gradient method employed a linear gradient ramping from 3% B to 36% B in 5 minutes (Solvent A: 0.1% formic acid (FA); Solvent B: acetonitrile (ACN)/0.1% FA) with a flow rate of 500 μl/min. The column was washed by an increase to 80% B in 0.5 minutes, followed by the flow rate being increased to 850 μl/min and the system kept at this setting for 0.2 minutes. In the next 0.1 minutes the B proportion was reduced to 3%, the flow rate was then reduced to 600 μl/min in 1.2 minutes and further to 500 μl/min in 0.5 min, the column was then equilibrated for 0.3 min. The 3-minute one-column active gradient method started with an equilibration for 0.1 min at 3% B and 500 μl/min flow rate and then employed a linear gradient ramping from 3% B to 32% B in 2.55 minutes followed by an increase to 40% B in 0.35 minutes. In the next 0.5 minutes the B proportion was increased to 80% and the flow rate to 850 μl/min, with the system kept at this setting for 0.2 minutes. In the next 0.1 minutes the B proportion was reduced to 3%, the flow rate was then reduced to 500 μl/min in 1.0 minute, and the column was equilibrated for 0.7 min.

The two-column system (1290 Infinity II (Agilent)) with two binary pumps connected to two positions of the ten port valve was equipped with two 30 × 2.1 mm Luna OMEGA 1.6 μm C18 100 Å (Phenomenex) columns operated at 60°C. The system was scheduled to use two sequential LC methods to work in alternating column regeneration mode. The pump 1 linear gradient ramped from 3% to 36% B in 3 min (850 μl/min), and simultaneously the pump 2 in the first 0.2 min was kept at 3% B (850 μl/min) with subsequent increase in 0.3 min to 80 % B and to 1000 μl/min. The flow rate was then increased to 1200 μl/min in the next 0.5 min and kept at that setting for the next 0.4 min. This wash was followed by a reduction to 3% B and the flow to 850 μl/min in 0.1 min, then equilibration for 1.5 min. Once the next method was started, after the valve switch, the same gradients were assigned to the opposite pumps, and the sample was injected into the washed and equilibrated column.

The positive m/z range was calibrated using four or five ions detected in the Agilent ESI-Low Tuning Mix (m/z [Th], 322.0481, 622.0289, 922.0097, 1221.990, and 1521.9714). For MS calibration in the ion mobility dimension, two ions were selected (m/z [Th], 1/*K*_0_: 622.0289, 0.9848; 922.0097, 1.1895). The selection of a third ion for ion mobility calibration is also possible.

### Peptide digests used during this study

MS-Compatible Human Protein Extract Digest (K562) was purchased from Promega (V6951), MassPREP *E. coli* Digest Standard was purchased from Waters (186003196). Human plasma digests prepared previously using a high-throughput robotics platform^5^ were used.

### Library generation

The K562 digest was fractionated using the High-pH Reverse-Phase Peptide Fractionation Kit (Pierce, 84868) according to the protocol provided by the manufacturer. In total 3 μg of isolated peptides for most of the fractions (except pH 7 and 8 for which 1.7 and 1.0 μg were used, respectively) were analyzed using a 30 × 2.1 mm Luna OMEGA 1.6 μm C18 100 Å (Phenomenex) column at 60°C using a linear gradient ramping from 3% B to 36% B in 20 minutes (Buffer A: 0.1% FA; Buffer B: ACN/0.1% FA) with a flow rate of 500 μl/min. In the next 0.5 min B was increased to 80% and in the next 0.1 min the flow was increased to 0.85 μl/min and kept at this setting for 0.2 min. In the next 0.1 min B was reduced to 3% and the flow was decreased to 600 μl/min after 1.2 min and to 0.50 μl/min after 0.1 min and kept at that setting for 0.3 min. The accumulation and ramp times were adjusted to 133 μs.

### Data analysis

The raw data were processed using DIA-NN 1.8.1^7,15^. Mass accuracies were set to 10 ppm for spectral library creation and to 15 ppm for all other analyses. Scan window was set to 0 (automatic) for spectral library creation and plasma dataset analysis and was set to 6 for all other analyses. MBR was disabled for the analysis with a DIA-based spectral library and enabled for the two-species benchmark and the plasma dataset analysis. Library generation was set to “Smart profiling” for the plasma analysis and “IDs, RT & IM profiling” for other analyses. For the generation of the in silico predicted spectral libraries from sequence databases (human and *E. coli*), the precursor charge range was restricted to 2–3. The plasma analysis was performed using a public spectral library^24^, which had information in it replaced with in silico predicted using the “Deep learning-based spectra, RTs and IMs prediction” option in DIA-NN. All other settings were kept at default.

The precursor FDR threshold was set to 1% for all DIA-NN analyses. For protein-level analyses, the following protein group q-value filtering was applied: 1% run-specific for the analysis of the dilution series; 1% global for the analysis of the two-species benchmark; 1% global for the analysis of the plasma dataset. The analysis of the plasma data was performed at the gene group level, ‘protein’ numbers on all figures for this experiment thus refer to the numbers of gene groups.

The data post-processing analysis was performed in R version 3.14. In the dilution series analysis, out of multiple injections of the same amount, the first three were considered. Differential abundance analysis depending on COVID-19 disease status and severity was performed by applying the Kendall Tau trend test for the log_2_-transformed protein levels against the disease severity encoded as the WHO grade (0: healthy; 3–7: mild to critical), using the EnvStats 2.7.0 R package. P-values were adjusted via Benjamini-Hochberg multiple testing correction using the p.adjust() R function. Heatmap generation was performed using the ComplexHeatmap 2.10.0 R package.

## Author contributions

Conception: M.R., V.D., L.Sz.; experimental design: V.D. and L.Sz.; mass spectrometry: L.Sz.; data analysis: V.D., C.G., L.Sz.; supervision: M.R., V.D., M.M.; initial draft: V.D., M.R., L.Sz.; writing: all authors edited and approved the final manuscript.

## Acknowledgements

We thank Florian Kurth, Leif-Erik Sander, Martin Witzenrath, Stefan Hippenstiel, Christof von Kalle, Andreas Hocke, Wolfgang Kübler, and the PA-COVID-19 study group (all Charité – Universitätsmedizin Berlin) and their teams for a long-standing collaboration in the COVID-19 studies, specifically plasma proteomics. Further, we thank Daniela Ludwig, Agathe Niewienda, and Torsten Schwecke for their help with proteomic sample preparation. This work was supported by the German Ministry of Education and Research (BMBF), as part of the National Research Node “Mass spectrometry in Systems Medicine” (MSCoreSys), under grant agreements 031L0220A (to M.R.) and 161L0221 (to V.D.), and in part by the European Research Council (ERC) under grant agreement ERC-SyG-2020 951475 (to M.R). Figure 2a was created with BioRender.com.

## Supplementary figures

**Supplementary Figure S1.**
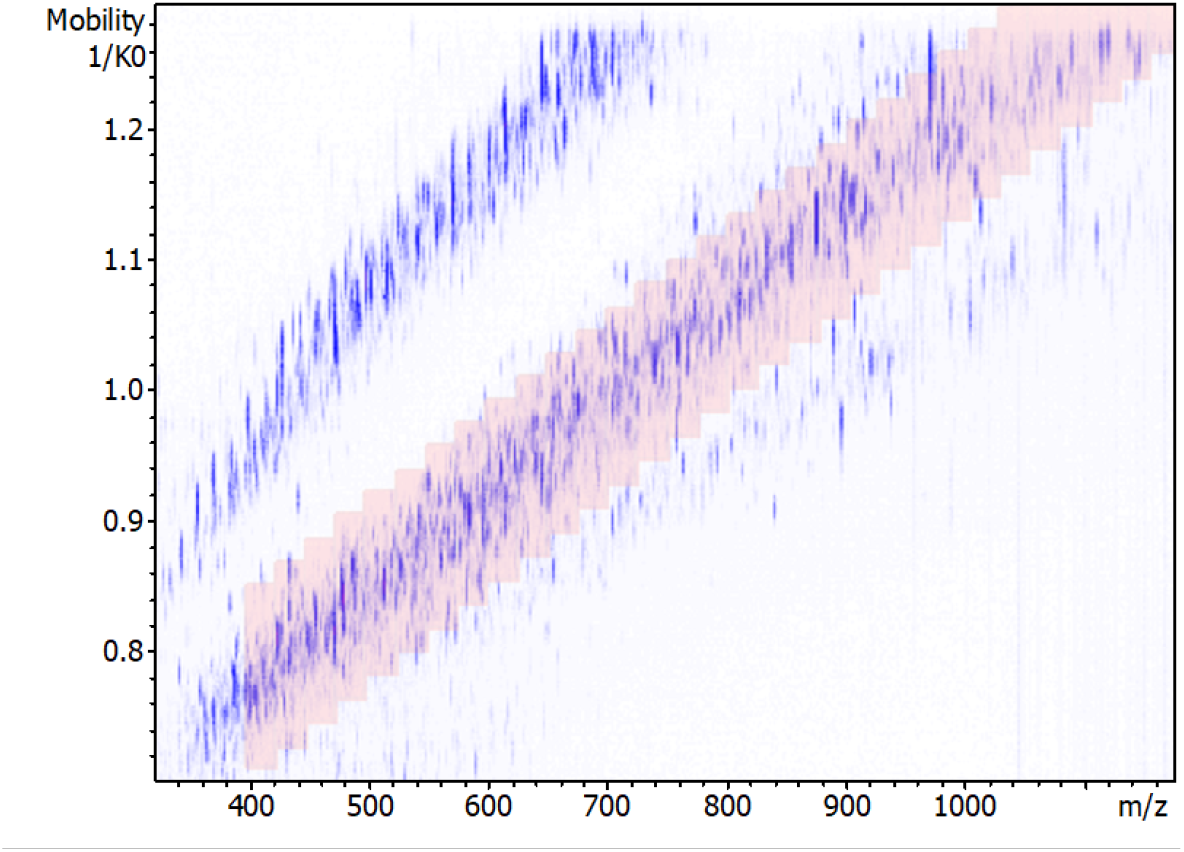
dia-PASEF acquisition scheme overlaid with the ion cloud for an injection of 2 μg of a K562 tryptic digest.

